# Conservation at the uterine-placental interface

**DOI:** 10.1101/2022.06.14.496152

**Authors:** Regan L. Scott, Ha T. H. Vu, Ashish Jain, Khursheed Iqbal, Geetu Tuteja, Michael J. Soares

## Abstract

The hemochorial placentation site is characterized by a dynamic interplay between trophoblast cells and maternal cells. These cells cooperate to establish an interface required for nutrient delivery to promote fetal growth. In the human, trophoblast cells penetrate deep into the uterus. This is not a consistent feature of hemochorial placentation and has hindered the establishment of suitable animal models. The rat represents an intriguing model for investigating hemochorial placentation with deep trophoblast cell invasion. In this study, we used single cell RNA sequencing to characterize the transcriptome of the invasive trophoblast cell lineage, as well as other cell populations within the rat uterine-placental interface during early (gestation day, **gd**, 15.5) and late (gd 19.5) stages of intrauterine trophoblast cell invasion. We identified a robust set of transcripts that define invasive trophoblast cells, as well as transcripts that distinguished endothelial, smooth muscle, natural killer, and macrophage cells. Invasive trophoblast, immune, and endothelial cell populations exhibited distinct spatial relationships within the uterine-placental interface. Furthermore, the maturation stage of invasive trophoblast cell development could be determined by assessing gestation-stage dependent changes in transcript expression. Finally, and most importantly, expression of a prominent subset of rat invasive trophoblast cell transcripts is conserved in the invasive extravillous trophoblast cell lineage of the human placenta. These findings provide foundational data to identify and interrogate key conserved regulatory mechanisms essential for development and function of an important compartment within the hemochorial placentation site that is essential for a healthy pregnancy.

**SIGNIFICANCE:** Trophoblast cell-guided restructuring of the uterus is an essential event in the establishment of the hemochorial placenta. Establishment of a suitable animal model for investigating regulatory mechanisms in this critical developmental process is a key to better understanding the etiology of diseases of placentation, such as early pregnancy loss, preeclampsia, intrauterine growth restriction, and preterm birth. The rat exhibits deep trophoblast cell invasion, as seen in human hemochorial placentation. Similarities are identified in the transcriptomes of rat and human invasive trophoblast cells, leading to the discovery of conserved candidate regulators of the invasive trophoblast cell lineage. This creates opportunities to test hypotheses underlying the pathophysiologic basis of trophoblast cell-guided uterine transformation and new insights into the etiology of diseases of placentation.

## INTRODUCTION

The uterine-placental interface is a dynamic site where trophoblast and uterine cells cooperate to support growth and maturation of the fetus. These tasks are accomplished by specialized trophoblast cells, which arise from a multi-lineage cell differentiation pathway (**1**). Among specialized trophoblast cells are those that acquire invasive properties and enter the uterine parenchyma where they facilitate the transformation of the uterine environment (**2-5**). In some primates and rodents, trophoblast cells penetrate the uterine vasculature and are directly bathed by maternal blood, a process referred to as hemochorial placentation (**6, 7**). Trophoblast cells entering the uterus are generically termed invasive trophoblast cells. In the human placenta, they are called extravillous trophoblast (**EVT**) cells. Invasive trophoblast cells migrate inside uterine blood vessels where they replace the endothelium (endovascular) and external to the vasculature where they move among uterine stromal cells (interstitial). Impairments in trophoblast-directed uterine transformation are linked to pregnancy-related diseases (**2, 3, 8**). However, there is a limited understanding of the regulatory mechanisms controlling the function of invasive trophoblast cells.

Interactions between invasive trophoblast cells and uterine cells are critical for the establishment of pregnancy and may be best investigated *in vivo*. Depth and extent of intrauterine trophoblast cell invasion show prominent species differences (**9**), affecting selection of suitable *in vivo* models. Human hemochorial placentation is characterized by deep intrauterine trophoblast cell invasion, a feature not shared with the mouse (**6**). In contrast to the mouse, the rat uterine-placental interface is characterized by deep intrauterine trophoblast cell invasion (**9-11**). This structural feature of the rat uterine-placental interface, and the availability of single cell RNA sequencing (**scRNA-seq**) technology, make the capture of invasive trophoblast cells remarkably straightforward.

In this report, scRNA-seq was performed on the rat uterine-placental interface to profile cell populations within the structure, and to identify conserved candidate regulators of invasive trophoblast cell lineage development.

## RESULTS

### Identification of cell clusters within the uterine-placental interface

Intrauterine trophoblast cell invasion in the rat is first detected at midgestation in the form of endovascular invasive trophoblast cells lining arterioles that penetrate the mesometrial decidua (**10**). After gd 13.5, trophoblast cells exit the junctional zone of the placenta penetrating beyond the decidua and deep into the mesometrial uterine parenchyma (**10, 16**). Both endovascular and interstitial invasive trophoblast cells are evident beginning on gd 14.5 (**10, 12**). The uterine-placental interface is the nodule of uterine tissue juxtaposed to the decidua and placenta and the destination of trophoblast cell invasion. This structure is retained in the uterus following removal of the placenta and adherent decidua and is easily dissected for further analysis (**Fig. 1A**) (**13**). The uterine-placental interface consists of an assortment of cell types, including invasive trophoblast, endothelial, immune, stromal, and smooth muscle cells. We performed scRNA-seq of the uterine-placental interface on gd 15.5 and 19.5, which reflect early and mature stages of intrauterine trophoblast cell invasion, respectively.

**Figure 1.**
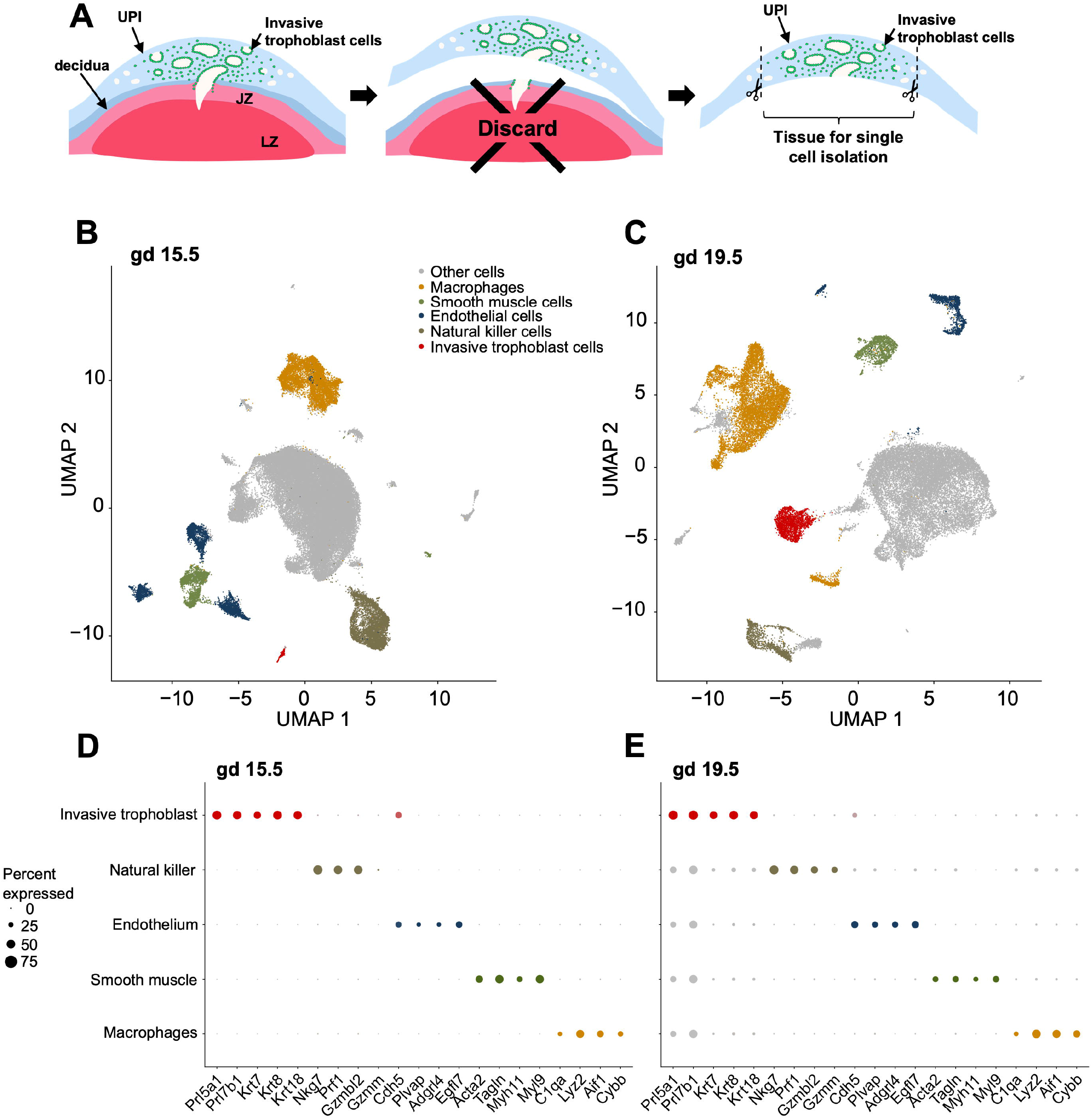
Single cell interrogation of the rat uterine-placental interface. **A**) Schematic showing isolation of the rat uterine-placental interface from gestation days (**gd**) 15.5 or 19.5. Uterine tissue at the site of trophoblast invasion was peeled away from placental tissue and associated decidua. **B, C**) Visualization of cell clusters from gd 15.5 and 19.5, respectively, plotted using uniform manifold approximation and projection (**UMAP**). **D, E**) Dot plots for gd 15.5 and 19.5 showing marker transcripts used to identify cell clusters. Dot size represents the average percentage of cells expressing the transcript. Dot colors correspond to cell types. For transcript expression level, see ***SI Appendix*, Fig. S2. and Dataset S1**. Abbreviations: JZ, junctional zone; LZ, labyrinth zone.

Following sequencing and data quality control (***SI Appendix*, Fig. S1)**, there were 65,842 cells in gd 15.5 samples, and 33,617 cells in gd 19.5 (***SI Appendix*, Fig. S2**). Cell clusters were then defined based on their transcript profiles (see Materials and Methods; **Fig. 1B, C; Dataset S1**). Endothelial, macrophage, natural killer (**NK**), invasive trophoblast, and smooth muscle cell clusters were identified using established biomarkers for the cell types (**Fig. 1D, E; *SI Appendix*, Fig. S2**). Invasive trophoblast cells were conspicuous in their epithelial character (e.g. expression of cytokeratins: *Krt7, Krt8, Krt18*) and expression of members of the expanded prolactin (**PRL**) gene family (*Prl7b1, Prl5a1, Prl4a1, Prl2a1, Prl2a1, Prl5a2, Prl2c1, Prl6a1*) (**Dataset S1**). To date, expression of cytokeratin and a subset of PRL gene family members have formed the basis of tracking invasive trophoblast cells within the rat uterine-placental interface (**10, 14-21**).

### Spatial relationships of invasive trophoblast cells and uterine cell populations

In situ hybridization was used to investigate spatial relationships of the uterine-placental interface cellular constituents at gd 15.5 and 19.5. Probes for identifying the cell types were selected based on the scRNA-seq cluster profiles (**Fig. 2A**). Invasive trophoblast cells were readily identified by the expression of the epithelial cell marker, *Krt8* and by expression of the invasive trophoblast cell marker, *Prl7b1* (**10, 14**), which were co-localized to both endovascular and interstitial trophoblast cells within the uterine-placental interface (**Fig. 2B, *SI Appendix, Fig S3***). Distributions of other transcripts, enriched in the invasive trophoblast cell cluster, within the rat placentation site were tracked by co-localization with *Prl7b1* (**Fig. 2C**). Although we were not able to resolve subpopulations of invasive trophoblast cells through scRNA-seq analysis, we were able to able to define distinct expression profiles for specific transcripts enriched in the invasive trophoblast cell cluster, which included expression patterns within the placentation site and among invasive trophoblast cells.

**Figure 2.**
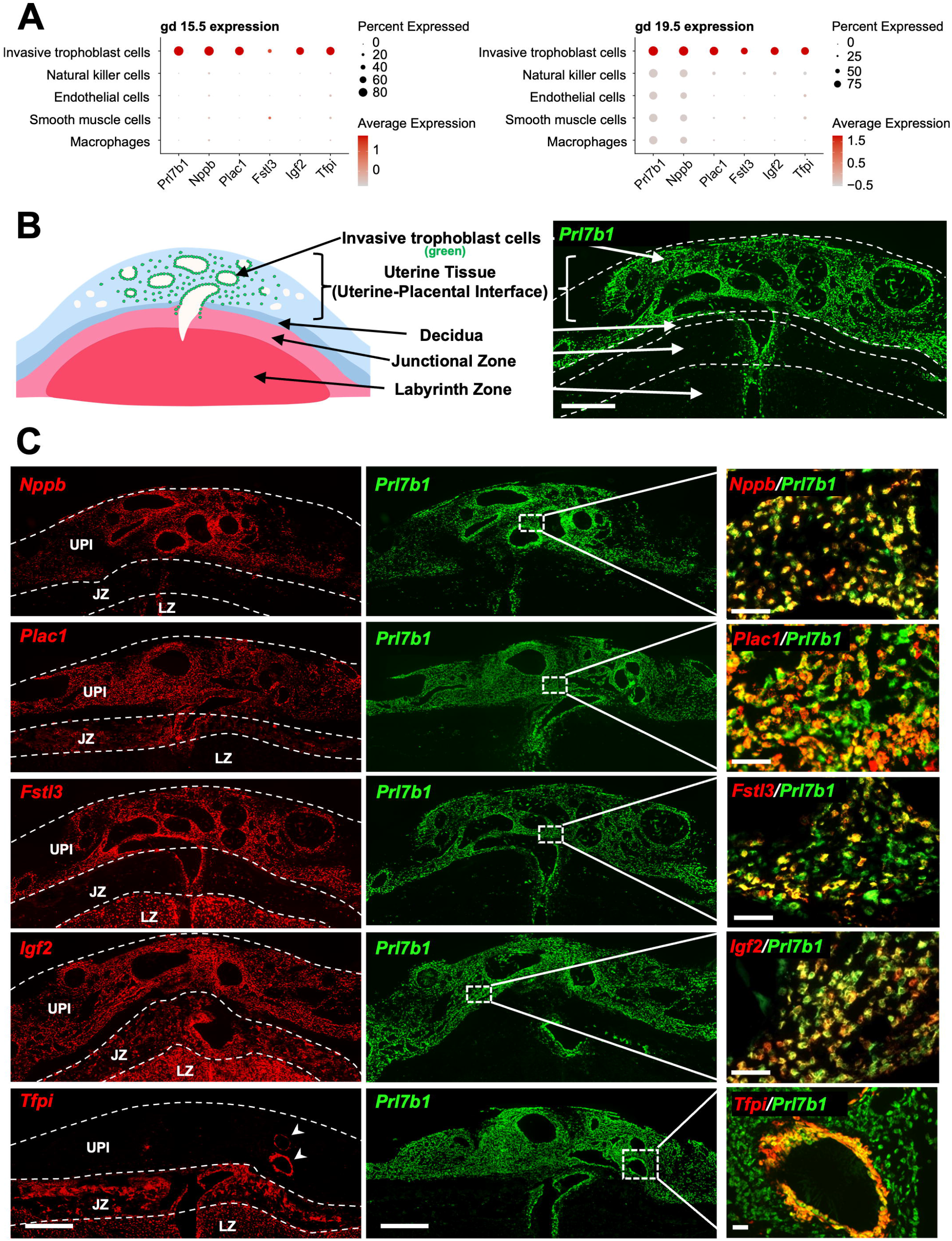
Distribution of transcripts enriched in invasive trophoblast cells at the rat uterine-placental interface (UPI). **A**) Dot plots for gestation day (gd) 15.5 (left panel) and 19.5 (right panel) showing expression levels of invasive trophoblast cell enriched transcripts. Dot size and color represents the average percentage of cells expressing the transcript and the average level of expression, respectively. **B)** Schematic depicting a gd 19.5 placentation site and a section through central region of a gd 19.5 placentation site. Distribution of invasive trophoblast cells was determined by in situ hybridization of *Prl7b1* transcripts. Scale bar=1000 µm. **C**) Localization of invasive trophoblast cell enriched transcripts [*Nppb, Plac1* (*LOC102550080*), *Fstl3, Igf2, Tfpi*; red] within the gd 19.5 placentation site. Transcripts were detected using in situ hybridization and co-localized to the distribution of *Prl7b1* (green). Scale bar (left and middle panels) = 1000 µm, scale bar (right panels) = 100 µm. Abbreviations: UPI, uterine-placental interface; JZ, junctional zone; LZ, labyrinth zone.

A range of placentation site associated patterns were observed. *Prl7b1* and *Nppb* were notable in their enrichment in only invasive trophoblast cells (**Fig. 2C**). *Plac1* (LOC102550080) was expressed in both invasive trophoblast cells and the junctional zone, whereas *Fstl3* was enriched in invasive trophoblast cells and the labyrinth zone (**Fig. 2C**). *Igf2* and *Tfpi* exhibited expression throughout the placentation site (**Fig. 2C**). Among invasive trophoblast cell enriched transcripts, *Prl7b1, Igf2*, and *Nppb* showed similar distributions in both endovascular and interstitial trophoblast cells (**Fig. 2C**). *Tfpi* expression was enriched in endovascular trophoblast cells (**Fig. 2C**), whereas *Plac1* was prominently expressed in interstitial trophoblast cells and a subset of endovascular trophoblast cells (**Fig. 2C**). *Fstl3* was localized to subsets of endovascular and interstitial trophoblast cells (**Fig. 2C**). The expression patterns are intriguing and imply that there are subpopulations of invasive trophoblast cells, which were not resolved by scRNA-seq.

Differences were also identified in the spatial relationships of invasive trophoblast cells with immune cells. Invasive trophoblast cells (*Prl7b1* positive) and NK cells (*Prf1* positive) exhibited a reciprocal relationship and showed little overlap in their spatial distribution (**Fig. 3A**), while macrophage populations (*Lyz2* positive) were interspersed among invasive trophoblast cells within the uterine placental interface (**Fig. 3B**). In situ hybridization of endothelial cell cluster transcripts indicated the existence of subsets of endothelial cells, including the presence of peripherally located blood vessels in late gestation placentation sites that were not restructured by endovascular invasive trophoblast cells (***SI Appendix*, Fig. S4**). *Adgrl4* and *Cdh5* showed some evidence for dual expression in endothelial cells and endovascular trophoblast cells, while *Cdh5* also exhibited expression in interstitial invasive trophoblast cells (***SI Appendix*, Figs. S4 and S5**).

**Figure 3.**
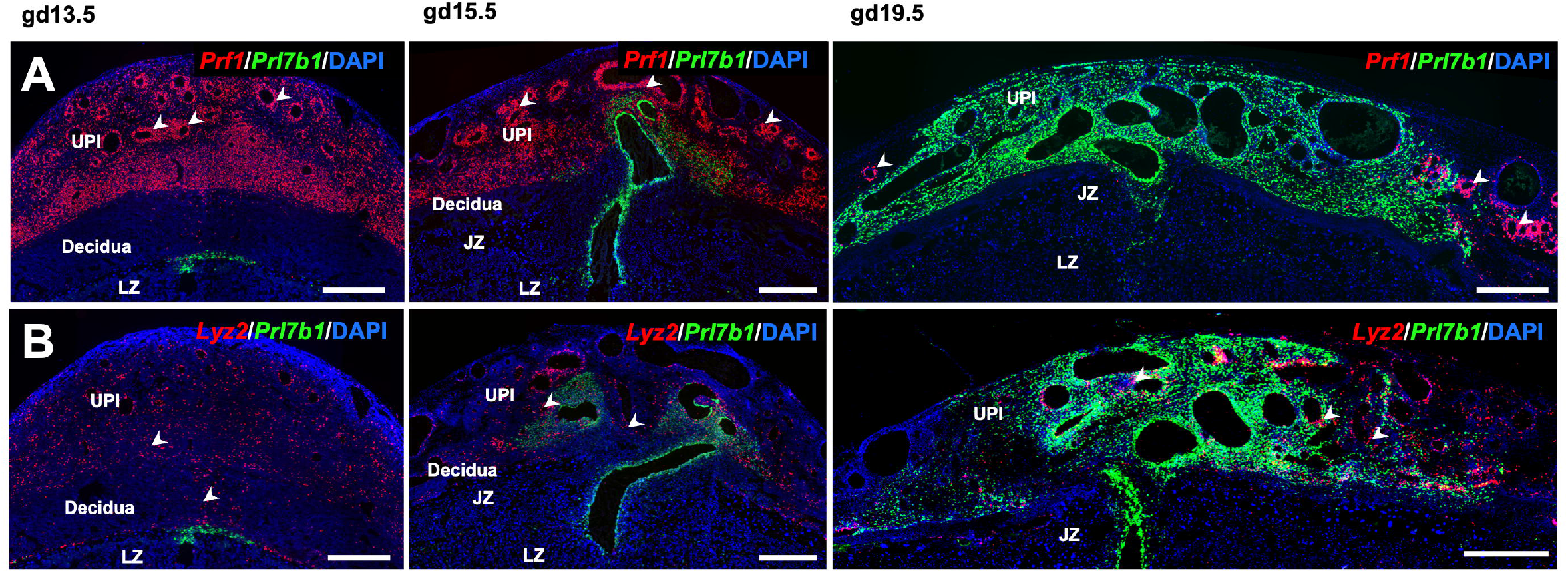
Distribution of NK cells and macrophages within the rat uterine-placental interface. **A**) NK cell and invasive trophoblast cells were monitored in gd 13.5, 15.5, and 19.5 placentation sites using in situ hybridization for *Prf1* (red) and *Prl7b1* (green), respectively. **B**) Macrophages and invasive trophoblast cells were monitored in gd 13.5, 15.5, and 19.5 placentation sites using in situ hybridization for *Lyz2* (red) and *Prl7b1* (green), respectively. Scale bar=1000 µm. Arrowheads indicate examples of the distribution of NK cells (top panels, *Prf1* positive) and macrophages (bottom panels, *Lyz2* positive). Abbreviations: UPI, uterine-placental interface; JZ, junctional zone; LZ, labyrinth zone.

### Gestation stage-dependent invasive trophoblast cell gene expression

Differential patterns of transcript expression were observed for invasive trophoblast cells during the initial phase of intrauterine invasion (gd 15.5: 115 enriched transcripts) versus late-stage invasion (gd 19.5: 126 transcripts) (**Fig. 4A, Dataset S2**). To explore the biological functions associated with the differentially expressed transcripts, we carried out gene ontology (**GO**) analysis. GO terms related to protein translation (“cytoplasmic translation”, -log_10_(q-value) = 2.49) and detoxification (“detoxification”, - log_10_(q-value) = 1.78) were significantly enriched with transcripts upregulated at gd 15.5, whereas transcripts upregulated at gd 19.5 were significantly linked to female pregnancy (“female pregnancy”, -log_10_(q-value) = 15.23) and epithelial cell migration (“positive regulation of epithelial cell migration”, -log_10_(q-value) = 2.54) (**Fig. 4B, Dataset S2**). Some of the top transcripts differentially expressed at gd 15.5 and gd 19.5 by invasive trophoblast cells were unique to rodents (gd 15.5: *LOC171573, Ceacam9, Doxl1*; gd 19.5: *Prl7b1, LOC684107*; **Fig. 4A**). *LOC171573* and *Ceacam9* were conspicuous in the initial wave of invasive trophoblast cells penetrating into the uterus (**Fig. 4C, D**). The biology of LOC171573, also called spleen protein 1 precursor, and CEACAM9 in placentation is not well understood (**22, 23**). CEACAM9 is a member of the larger carcinoembryonic antigen family, which has been implicated in cell adhesion, epithelial barrier function, and inflammatory responses (**24**). An implication from these observations may be that invasive trophoblast cells first entering deep into the potentially hostile uterus activate species-specific protective mechanisms, which are not required later in gestation as the uterine parenchyma becomes conditioned by infiltrating invasive trophoblast cells.

**Figure 4.**
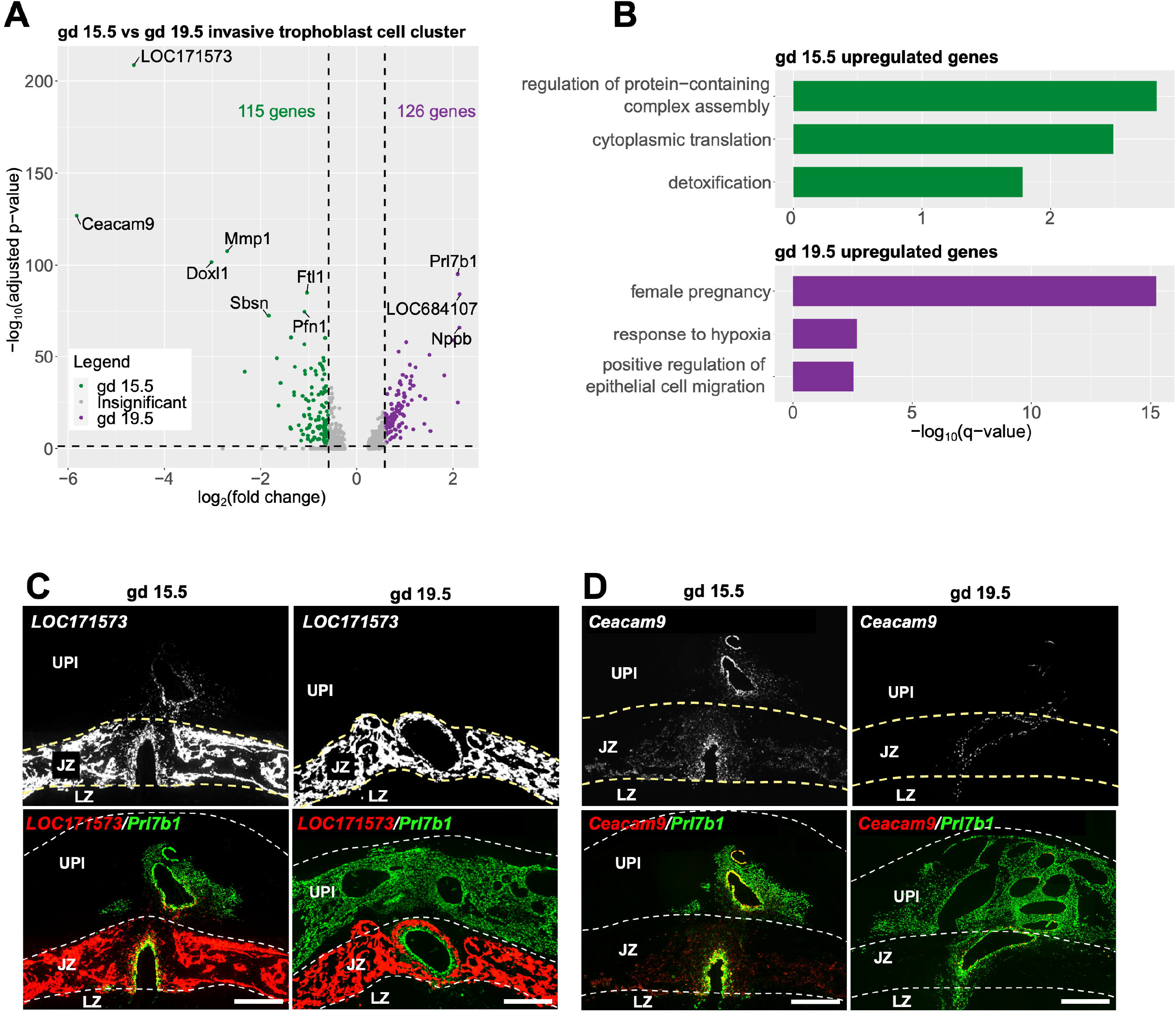
Differential invasive trophoblast cell gene expression on gestation day (gd) 15.5 versus gd 19.5. **A**) Volcano plot showing differentially expressed genes (**DEGs**) in gd 15.5 and 19.5 invasive trophoblast cell clusters. Top 10 most significantly changed genes are labeled. **B**) Bar plots showing selected significantly enriched gene ontology terms for transcripts upregulated at gd 15.5 (top) and transcripts upregulated at gd 19.5 (bottom). For full list of enriched terms, see ***SI Appendix*, Dataset S2, panels C, D**) Differential expression of *LOC171573* and *Cecam9* transcripts within placentation sites at gd 15.5 and gd 19.5. Transcripts were detected using in situ hybridization and co-localized to the distribution of *Prl7b1* (green). Yellow arrows indicate the intrauterine invasive cells expressing *LOC171573* and *Ceacam9* at gd 15.5. Scale bar=1000 µm. Abbreviations: UPI, uterine-placental interface; JZ, junctional zone; LZ, labyrinth zone.

### Conservation of rat and human invasive trophoblast cell-specific transcripts

To explore invasive trophoblast cell transcripts conserved in rats and humans, we started by using the PlacentaCellEnrich webtool (**25**). Briefly, PlacentaCellEnrich groups genes that have cell-type specific expression according to scRNA-seq data generated in human placenta and then calculates cell-type-specific enrichment for a given set of input genes (e.g., invasive trophoblast cell cluster marker genes). PlacentaCellEnrich employs the definitions of cell-type specific gene groups from the Human Protein Atlas (**26**), in which a gene can be cell-type specific if it is highly expressed in a group of cell types compared to the rest of the cell types. Therefore, a single gene can be classified as having cell-type specific expression in multiple trophoblast cell types. Interestingly, we observed that transcripts marking the invasive trophoblast clusters at gd 15.5 and gd 19.5 were most significantly enriched for EVT cell-specific genes when using cell-type-specific groups defined through five independent single cell human placenta or early embryo culture datasets **(Fig. 5A) (27-31)**.

**Figure 5.**
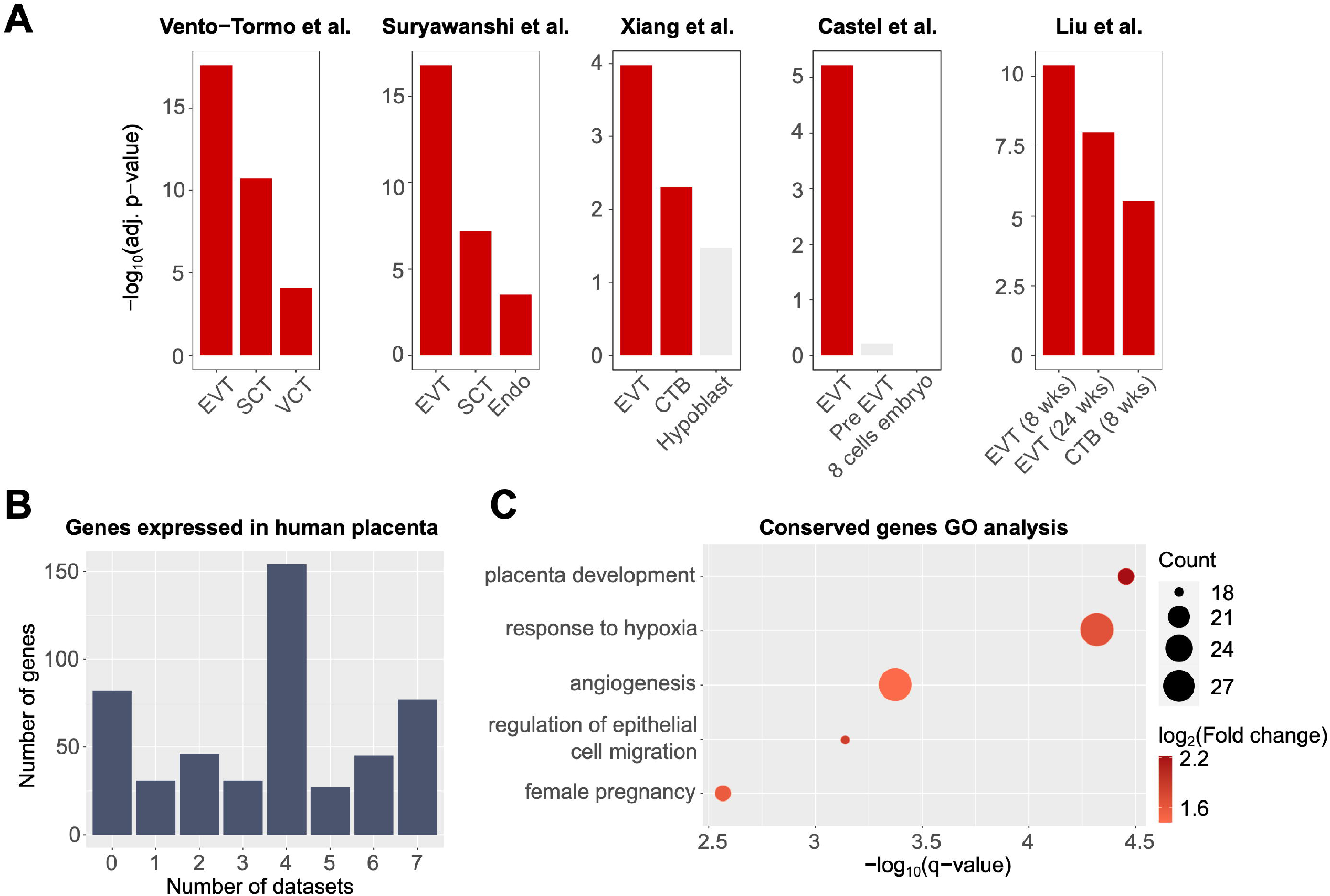
Conserved transcript expression in rat invasive trophoblast cells and human extravillous trophoblast (EVT) cells. **A**) Bar plots showing that transcripts highly expressed in the invasive trophoblast cell clusters at gd 15.5 and 19.5 in rats share similar profiles to EVT cells in human placenta. Enrichment analyses were carried out using human placenta single cell data as references: Vento-Tormo *et al*. (**29**), Suryawanshi *et al*. (**28**), Xiang *et al*. (**30**), Castel *et al*. (**31**), and Liu *et al*. (**27**). A significant enrichment has an adjusted p-value ≤ 0.05, fold change ≥ 1.5, and number of observed genes ≥ 5. Red, significant enrichment; grey, insignificant enrichment. **B**) Bar plot showing the number of rat invasive trophoblast cell genes that were expressed in human placenta datasets. **C**) Dot plots showing gene ontology (**GO**) enrichment for rat invasive trophoblast cell genes that are also expressed in human placenta. Dot size represents the number of observed genes related to a term; dot colors correspond to log_2_(Fold change) of the terms. Selected gene ontology terms are shown. For a full list of enriched terms, see ***SI Appendix*, Fig. S6**. Abbreviations: EVT, extravillous trophoblast; SCT, syncytiotrophoblast; VCT, villous cytotrophoblast; Endo, vascular endothelial cells (villi); CTB, cytotrophoblast; EVT (8 wks), EVT from villi at 8 weeks of pregnancy; EVT (24 wks), EVT from decidua at 24 weeks of pregnancy; CTB (8 wks), CTB from villi at 8 weeks of pregnancy.

Motivated by the results from PlacentaCellEnrich, we further annotated all 493 of the invasive trophoblast cell cluster transcripts in rat at either gd 15.5 or gd 19.5 that have expressed human orthologs (see Materials and Methods) in not only single cell data but also bulk RNA-seq data obtained from EVT cells differentiated from human trophoblast stem cells (**32**) and EVT cells isolated from first trimester human placenta (**33**). The rat invasive trophoblast cell cluster transcripts showed some evidence of species-specificity as there were 82 (16.63%) transcripts that did not have human orthologs or were not expressed in human **(Figure 5B, *SI Appendix*, Table S1)**. However, the vast majority of transcripts (411, 83.37%) from the rat invasive trophoblast scRNA-seq cell clusters were also expressed in at least one of the human EVT datasets **(Figure 5B, *SI Appendix*, Table S1)**. Next, we used GO analysis on invasive trophoblast cell genes expressed in at least one dataset and observed that the genes are enriched for processes related to low oxygen conditions (“response to hypoxia”, -log_10_(q-value) = 4.32), pregnancy and placenta development (“female pregnancy”, -log_10_(q-value) = 2.57; “placenta development”, -log_10_(q-value) = 4.45), cell migration (“regulation of epithelial cell migration”, -log_10_(q-value) = 3.14), and vasculature development (“angiogenesis”, -log_10_(q-value) = 3.37) **(Figure 5C, *SI Appendix*, Table S2)**. Collectively, these observations reinforce the merits of modeling trophoblast-guided uterine transformation in the rat.

## DISCUSSION

Trophoblast cell-guided transformation of the uterus is a defining feature of the hemochorial placenta (**6, 7**). A specialized lineage of trophoblast cells referred to as invasive trophoblast or EVT cells executes these vital functions (**5, 6**). This fundamental process includes restructuring uterine spiral arteries to optimize blood flow into the placenta (**3, 5**). In human pregnancies, failures in trophoblast cell-guided uterine vasculature remodeling underly disease states such as preeclampsia, intrauterine growth restriction, and preterm birth (**8**). There is a need for modeling this key trophoblast cell-uterine interaction *in vivo*; however, arguments have been levied that the process as it occurs in human placentation is unique (**34**). Unfortunately, this opinion has stifled progress in understanding regulatory mechanisms controlling deep placentation. In this report, compelling evidence is provided supporting the utility of the rat as a model for investigating the uterine-placental interface. Two important characteristics of the rat placentation site are key features demonstrating its efficacy as a model: i) deep intrauterine trophoblast cell invasion (**9, 11, 35**) and ii) physical separation of intrauterine invasive trophoblast cells from the placenta (**36**). Cellular constituents of the rat uterine-placental interface were defined and are consistent with previous scRNA-seq analyses of the first trimester human uterine-placental interface (**27-29**). Furthermore, core conservation in the transcriptomes of rat and human invasive trophoblast/EVT cells was revealed.

There are two main populations of invasive trophoblast cells: i) interstitial and ii) endovascular. These two invasive trophoblast cell types exhibit positional and functional differences and are characteristic of both rat and human placentation sites (**3, 5, 9, 11, 35**). Invasive trophoblast cells and EVT cells did not resolve into clusters reflecting the two invasive trophoblast cell types in the present analysis or prior scRNA-seq efforts of the first trimester human placentation site (**27-29**). Spatial transcriptomic differences of interstitial versus endovascular trophoblast cells were resolved by in situ hybridization. This is especially noteworthy for *Tfpi* and *Mmp12* transcripts, which reside within the singular invasive trophoblast cell cluster, but based on in situ hybridization are enriched in endovascular trophoblast cells (**19, 37**). The reason for this discrepancy is not understood. It may relate to unequal recovery of interstitial versus endovascular trophoblast cells during the single cell isolation process. Alternatively, it may reflect the plasticity of the isolated invasive trophoblast cells and the requirement of positional cues for displaying phenotypic differences, which disappear once the cells are dissociated. Application of genome-wide spatial transcriptomic analyses (**38**) could further define phenotypic differences between interstitial and endovascular trophoblast cells.

Conservation in gene regulatory networks controlling rat and human invasive trophoblast and EVT cell lineages has been previously demonstrated and include *Mmp12, Ascl2*, and *Tfpi* (**19, 21, 37**). scRNA-seq analysis has expanded the list of candidate conserved regulators. Among these candidates are transcripts encoding transcriptional regulators (e.g. PPARG, CITED2), regulatory hubs in signaling pathways (e.g. IGF2, CDKN1C, FSTL3), and proteins known to directly affect cell migration and invasion (e.g. ITGA5, MMP12). Strategies exist in the rat for testing *in vivo* roles of candidates using global and conditional genome editing (**39**) and trophoblast cell-specific lentiviral mediated gene manipulation (**40, 41**). Dysregulation of candidate genes may directly affect the development and function of the invasive trophoblast cell lineage and indirectly affect uterine cell dynamics at the uterine-placental interface. These approaches can be further leveraged through comparative analysis *in vitro* using rat and human trophoblast stem cells (**32, 42**). In addition to conservation at the uterine placental interface there are also elements of species specificity, which is exemplified by the hormone producing capacity of invasive trophoblast and EVT cells (e.g. rat: prolactin family; human: chorionic gonadotropin and pregnancy specific glycoproteins).

In summary, conservation exists within the invasive trophoblast cell lineages of the rat and human providing a rationale for using the rat as an experimental model to elucidate regulatory mechanisms controlling uterine-placental cellular dynamics.

## MATERIALS AND METHODS

### Animals and tissue collection

Holtzman Sprague-Dawley rats were obtained from Envigo (Indianpolis, IN). Rats were housed on a 14 h light/10 h dark photoperiod with free access to food and water. Timed pregnancies were obtained by mating adult males (>10 weeks) and females (8-12 weeks). Pregnancies were confirmed with a sperm positive vaginal lavage, defined as gd 0.5. Uterine-placental interface tissues of the rat placentation site (also termed the metrial gland) from gd 15.5 and 19.5 were dissected as previously described (**36**) and used for scRNA-seq. Uteroplacental/fetal sites were also frozen in dry-ice cooled heptane and used for in situ hybridization localization of transcripts. Protocols for research with animals were approved by The University of Kansas Medical Center **(KUMC**) Animal Care and Use Committee.

### Cell isolation and scRNA-seq

Uterine-placental interface tissues were harvested from gd 15.5 (n=3 pregnancies/gd) and 19.5 (n=4 pregnancies/gd) rat placentation sites and placed in ice cold Hank’s Balanced Salt Solution (**HBSS**). Tissues were minced into small pieces with a razor blade and digested with Dispase II (1.25 units/ml, D4693, Sigma-Aldrich, St. Louis, MO), 0.4 mg/ml collagenase IV (C5138, Sigma-Aldrich), and DNase I (80 units/ml, D4513, Sigma-Aldrich) in HBSS for 30 min. Red blood cells were lysed using Ammonium-Chloride-Potassium Lysis buffer (A10492-01, Thermo-Fisher, Waltham, MA) with rotation at room temperature for 5 min. Intact cells were washed with HBSS supplemented with 2% fetal bovine serum (Thermo-Fisher), and DNase1 (Sigma-Aldrich) and passed through a 100 µm cell strainer (100ICS, Midwest Scientific, Valley Park, MO). Following enzymatic digestion, cellular debris was removed using MACS Debris Removal Solution (130-109-398, Miltenyi Biotec, Sunnyvale, CA). Cells were then filtered through a 40 µm cell strainer (40ICS, Midwest Scientific) and cell viability assessed, which ranged from 90-93%. Cells were used for the preparation of single-cell libraries using the 10X Genomics Chromium system (10X Genomics, Pleasanton, CA) and sequenced with an Illumina NovaSeq 6000 (Illumina, San Diego, CA) by the KUMC Genome Sequencing Facility (Kansas City, KS).

### scRNA-seq data analysis

*scRNA-seq preprocessing*. Raw data was aligned to the rat genome (Rnor 6.0, Ensembl 98) (**43**) and quantified using the Cell Ranger Software (version 4.0.0). The R package Seurat (version 4.1.0) (**44**) was used for quality control and downstream analysis. Hereinafter, if not mentioned, all parameter settings are as default. We retained the cells with the number of unique genes between 500 and 3500, and less than 20% of mitochondrial genes (***SI Appendix*, Fig. S1**). Next, we merged the replicates of gd 15.5 together using Seurat function Merge(). Since replicates of gd 19.5 were generated in two different batches, Seurat functions FindIntegrationAnchors() and IntegrateData() were used to perform batch correction and integrate gd 19.5 samples (***SI Appendix*, Fig. S1**). All mitochondrial genes were then removed from the resulting Seurat objects. To mitigate technical noise, we normalized, selected top 2000 most variable transcripts in each time point, then scaled the data. Principal component analysis (**PCA**) was then carried out with the top 2000 variable transcripts to reduce dimensions of the data. We accessed the significance of top 100 principal components (**PC**) in each time point with Seurat functions JackStraw() and ElbowPlot(). The analyses showed that PC 72 was the last significant component in the gd 15.5 sample (p-value ≤ 0.05); hence, we retained the first 72 PCs in gd 15.5 for downstream analyses. Similarly, the first 77 PCs were kept for the gd 19.5 sample (***SI Appendix*, Fig. S2**).

#### scRNA-seq clustering and cluster annotation

With the retained significant PCs, we utilized K-nearest neighbor (**KNN**) graphs and the original Louvain algorithm (**45**), which are implemented through the Seurat functions FindNeighbors() and FindClusters(), to identify cell clusters in each time point (resolution = 0.8). The cell clusters were then visualized with Uniform Manifold Approximation and Projection (**UMAP**). To identify marker transcripts for each cell cluster at each gestation day, we used the Seurat function FindAllMarker() with Wilcoxon Rank Sum test, which compares gene expression in each cell cluster to the expression across all of the other cell clusters. Original assays of individual replicates were inputs for the FindAllMarker() differential expression tests. A transcript was considered a marker of one cell cluster if it was expressed in ≥ 10% of the cells, its adjusted p-value was ≤ 0.05 and its average fold change was ≥ log_2_(1.5). We annotated the cell clusters using the following cell type markers: *Prl5a1, Prl7b1, Krt7, Krt8* and *Krt18* (invasive trophoblast cells); *Nkg7, Prf1, Gzmm* and *Gzmbl2* (NK cells); *Cdh5, Plvap, Adgrl4* and *Egfl7* (endothelial cells); *C1qa, Lyz2, Aif1* and *Cybb* (macrophages); *Acta2, Myl9, Tagln* and *Myh11* (smooth muscle cells). A cell cluster was annotated to a cell identity if it had ≥ 75% of the previously identified cell type markers.

#### scRNA-seq comparison between timepoints

To compare the expression profiles of trophoblast clusters between the two gestation days, we carried out differential expression analysis. We used the Seurat function FindMarkers() with the original assays of the samples as inputs for the differential tests. A differentially expressed gene at a gestation day was one with adjusted p-value ≤ 0.05 and average fold change was ≥ log_2_(1.5).

#### Gene ontology enrichment analysis

To explore gene set functions, we carried out gene ontology (**GO**) enrichment analysis using the R package clusterProfiler (version 3.16.1) (**46**). Gene annotation was obtained from the R package org.Rn.eg.db (version 3.11.4) (**47**). A fold change for each term was calculated as GeneRatio/BgRatio. Non-redundant terms were then obtained using the simplify() function in clusterProfiler. A GO term was considered enriched if it was non-redundant, its q-value was ≤ 0.05, fold change was ≥ 2, and the number of observed genes was 2 5.

#### Identification of conserved trophoblast genes between human and rat

We carried out cell enrichment analysis using the PlacentaCellEnrich webtool (**25**). Cell type-specific genes were identified based on human placenta or embryo culture single cell data (**27-31**). The datasets from Suryawanshi *et al*. (**28**) and Vento-Tormo *et al*. (**29**) were used as supplied in PlacentaCellEnrich (**25**). The datasets from Xiang *et al*. (**30**) and Castel et al. (**31**) were obtained from Seetharam *et al*. (**48**). The data from Liu *et al*. (**27**) was reanalyzed to build a reference dataset with transcripts per million (TPM)-like expression values following the PlacentaCellEnrich procedure (**25**). An enrichment was considered significant if its adjusted p-value was ≤ 0.05, fold change was ≥ 1.5 and the number of observed genes was ≥ 5. Code to carry out PlacentaCellEnrich analysis was adapted from the TissueEnrich R package (**49**). A gene was considered expressed in human EVT cells in Liu *et al*. (**27**), Suryawanshi *et al*. (**28**), Vento-Tormo *et al*. (**29**), Xiang *et al*. (**30**), and Castel *et al*. (**31**) datasets if its TPM-like expression was ≥ 1; expressed in Morey et al. human EVT cells (**33**) if its TPM is ≥ 1, and expressed in Okae et al. human EVT cells (**32**) if its FPKM (fragments per kilobase of TPM mapped reads) is ≥ 1.

#### *In situ* hybridization

Localization of transcript expression within placentation sites was performed on cryosections (20 µm) prepared from gd 15.5 and 19.5 rat placentation sites. RNAScope Multiplex Fluorescent V2 assay (Advanced Cell Diagnostics, Newark, CA) was used to detect transcripts within the uterine-placental interface according to protocols provided by the manufacturer. Probes were prepared to detect invasive trophoblast cells (*Krt8*: NM_199370.1, 873041-C2, target region: 134-1472; *Prl7b1*: NM_153738.1, 860181, 860181-C2, target region: 28-900; *Igf2*: NM_031511.2, 444561, target region: 1475-2634; *Fstl3*: NM_053629.3, 862961, target region: 2-766; *Nppb*: NM_031545.1, 583051, target region: 3-531; *Plac1*: NM_001024894.1, 860141, target region: 3-944; *Tfpi*: NM_017200.1, 878371, target region: 2-1138; *LOC171573*: NM_138537.2, 1104761, target region: 2-576); *Ceacam9*: NM_053919.2, 1166771, target region: 130-847); endothelial cell (*Adgrl4*: NM_022294.1, 878411-C2, target region: 886-1871; *Plvap*: NM_020086.1, 878401, target region: 61-1225), macrophage (*Lyz2*: NM_012771.3, 888811-C2, target region: 82-1181; natural killer cell (*Prf1*: NM_017330.2, 871601-C2, target region: 451-1452. A Nikon 80i upright microscope (Nikon, Melville, NY) and Photometrics CoolSNAP-ES monochrome was used to capture fluorescence images.

## Data and resource availability

scRNA-seq datasets are available at the Gene Expression Omnibus website (https://www.ncbi.nlm.nih.gov/geo/; GSE206086). All data generated and analyzed during this study are included in the published article and the online supporting files. All code used for the analyses are available at https://github.com/Tuteja-Lab/MetrialGland-scRNA-seq. Resources generated and analyzed during the current study are available from the corresponding author upon reasonable request.

## ACKNOWLEDGEMENTS

We thank Brandi Miller and Stacy Oxley for their assistance.

## AUTHOR CONTRIBUTIONS

R.L.S., H.T.H.V., G.T., and M.J.S. conceived and designed the research and wrote the manuscript; R.L.S., H.T.H.V., A.J. K.I., G.T., and M.J.S. performed experiments and/or analyzed data; all authors contributed to editing the manuscript.

## SI Appendix

**Figure S1.**
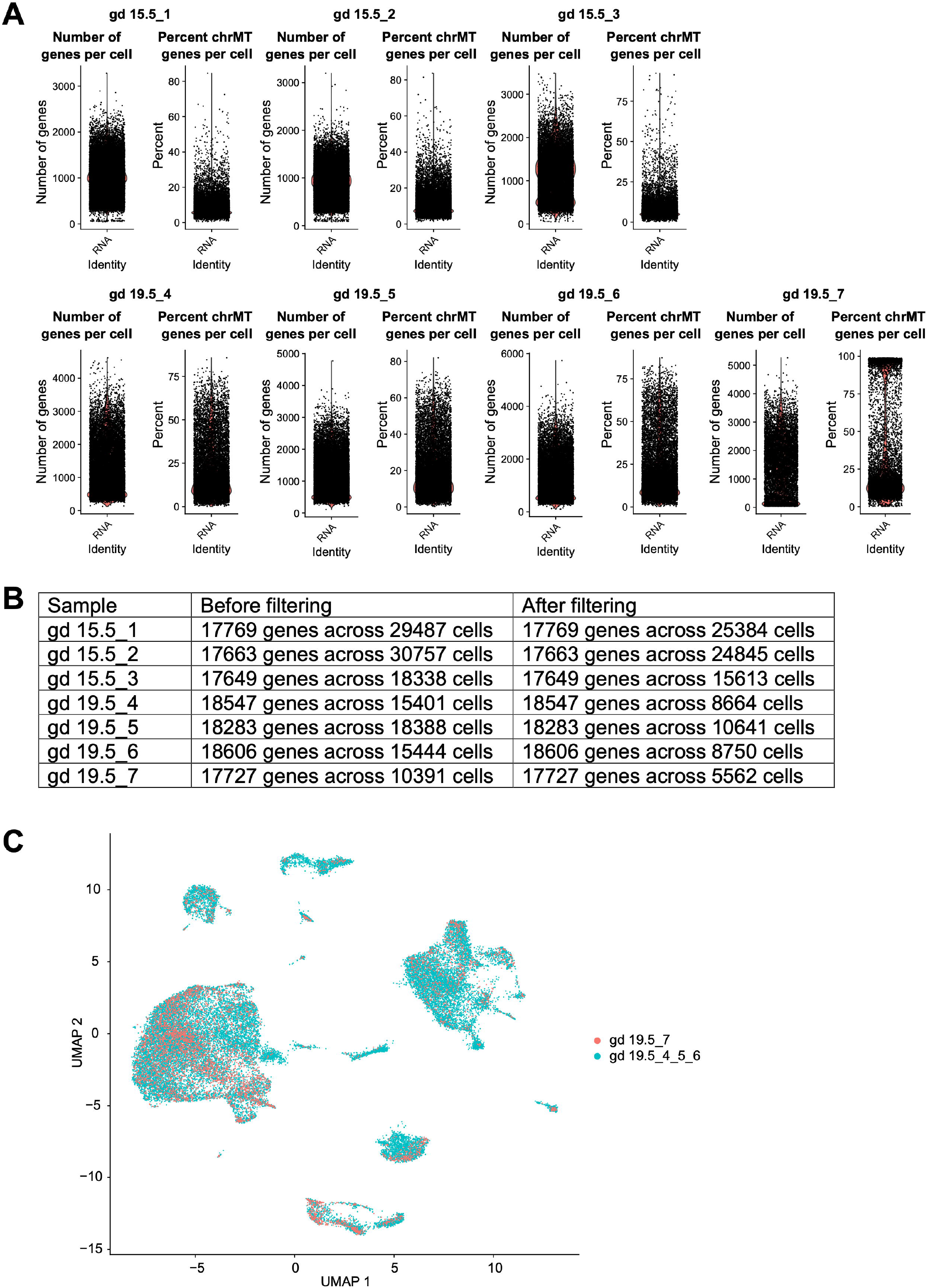
Quality control and processing of single cell RNA sequencing (scRNA-seq) data. **A**) Violin plots showing the distributions of number of genes and percentage of mitochondrial chromosome (chrMT) genes expressed per cell. Cells with the number of unique genes between 500 and 3500, and less than 20% of mitochondrial genes were retained. **B**) Summary table of the number of genes and cells retained in each sample after preprocessing. **C**) UMAP plot showing the cells from samples gd 19.5_4_ 5_6 and sample gd 19.5_7, which were generated in different batches, are integrated and well blended after the integration process.

**Figure S2.**
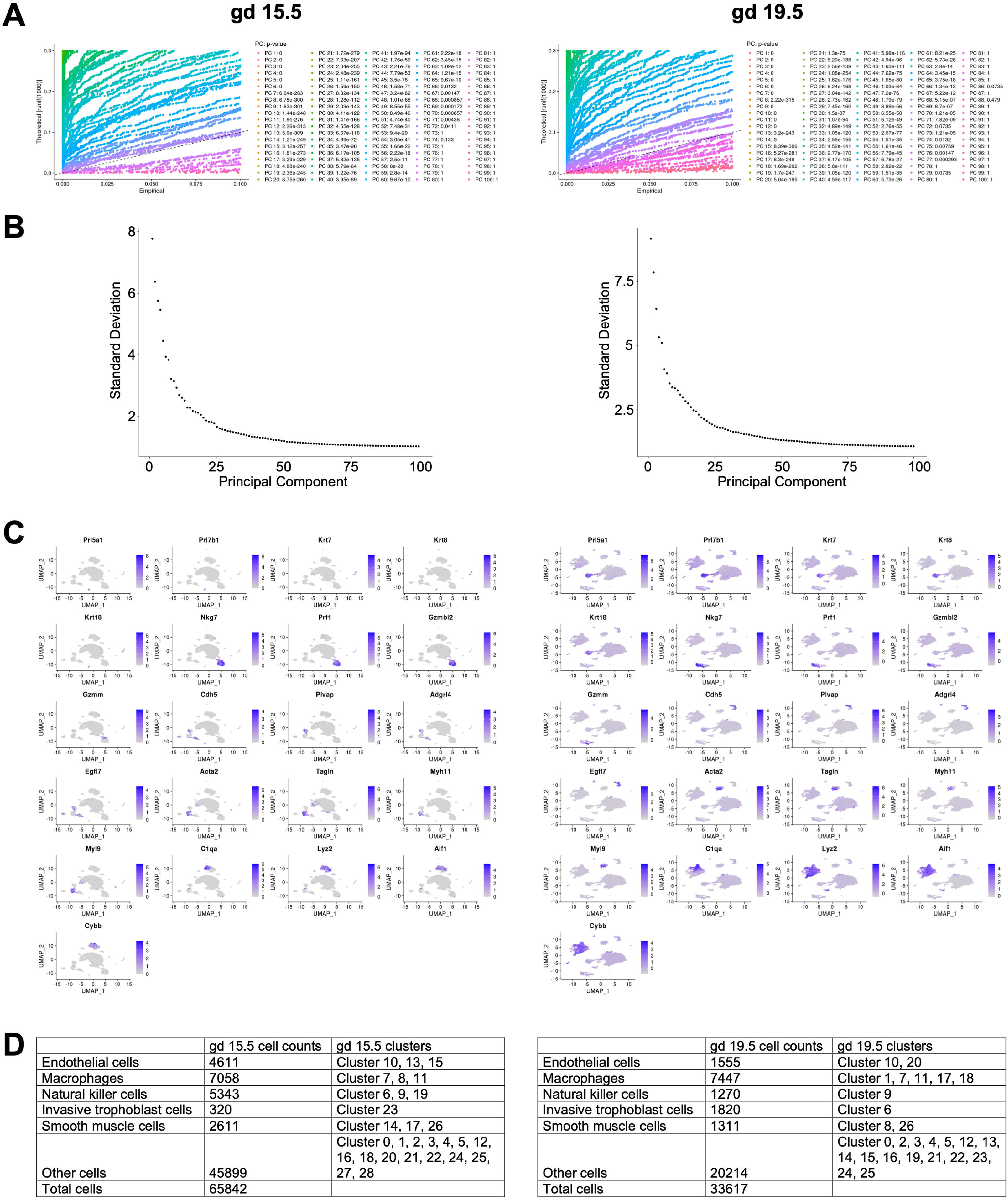
Clustering and cluster annotations. **A**) QQ plots showing the JackStraw analysis for principal component (**PC**) analysis significance. P-values indicating the significance of each PC were also shown. Based on the p-values, the first 72 PCs were used for the analyses on gd 15.5 samples, and the first 77 PCs were used for gd 19.5. Elbow plots showing the amount of standard deviation each PC represents. The first 72 PCs of gd 15.5 samples, and 77 PCs of gd 19.5 samples, captured the majority of the variation in the data. **C**) UMAP plots showing expression level of marker genes across cell clusters. **D**) Summary table of the cell cluster identities and number of cells in each cell group.

**Figure S3.**
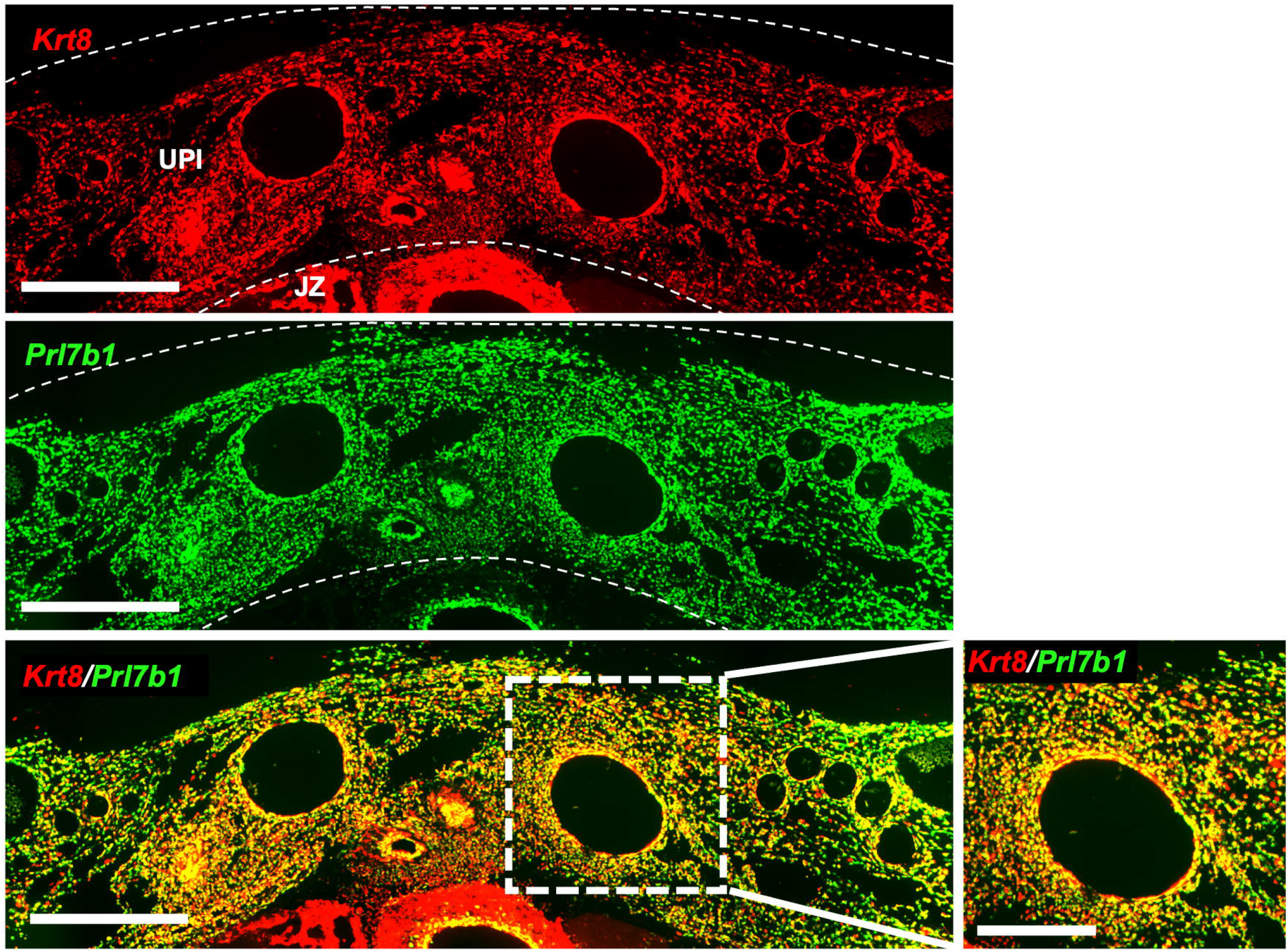
Colocalization of *Krt8* and *Prl7b1* in invasive trophoblast cells at the uterine-placental interface. In situ hybridization of gd 19.5 rat placentation site for *Krt8* (red; top panel), *Prl7b1* (green; center panel), and co-localization of *Krt8* and *Prl7b1* (bottom panels). Scale bar (left panels)=1000 µm, scale bar (right panel)=500 µm. Abbreviations: UPI, uterine-placental interface; JZ, junctional zone.

**Figure S4.**
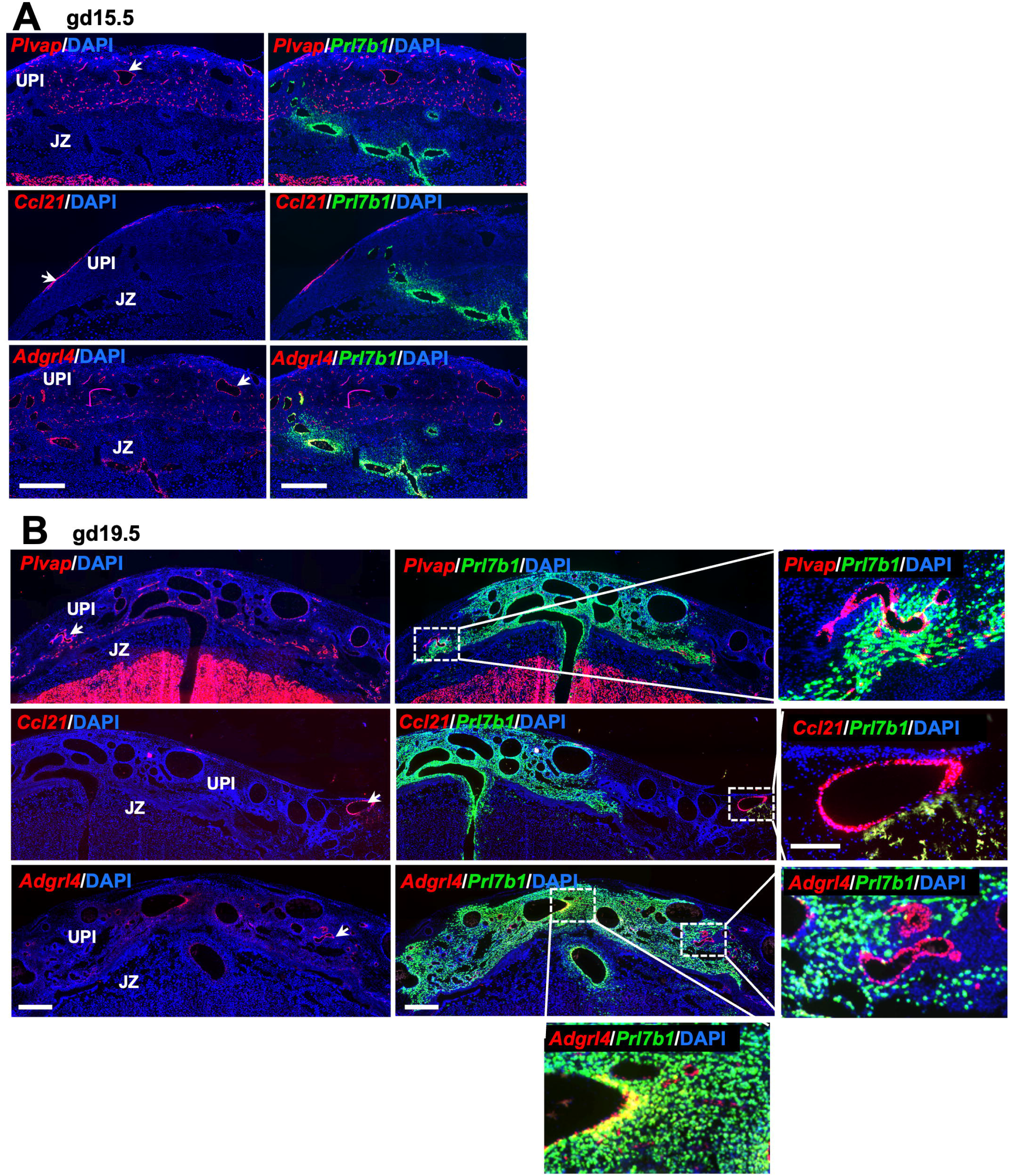
Endothelial cells at the uterine-placental interface. **A**) In situ hybridization of gd 15.5 rat placentation site for endothelial cell-specific transcripts *Plvap, Ccl21*, and *Adgrl4* (red; left panels) and localization of each endothelial cell marker with an invasive trophoblast cell-specific transcript (*Prl7b1*; right panels). Scale bar=1000 µm. **B**) In situ hybridization of gd 19.5 rat placentation sites for *Plvap, Ccl21, and Adgrl4*, and (red; left panels) and localization of each endothelial cell marker with an invasive trophoblast cell-specific transcript (*Prl7b1;* center, right, and bottom panels). Panels on the right and bottom are high magnification images of regions shown in the center panels. Scale bar (left and center panel)=1000 µm, scale bar (right and bottom panels)=500 µm. Abbreviations: UPI, uterine-placental interface; JZ, junctional zone.

**Figure S5.**
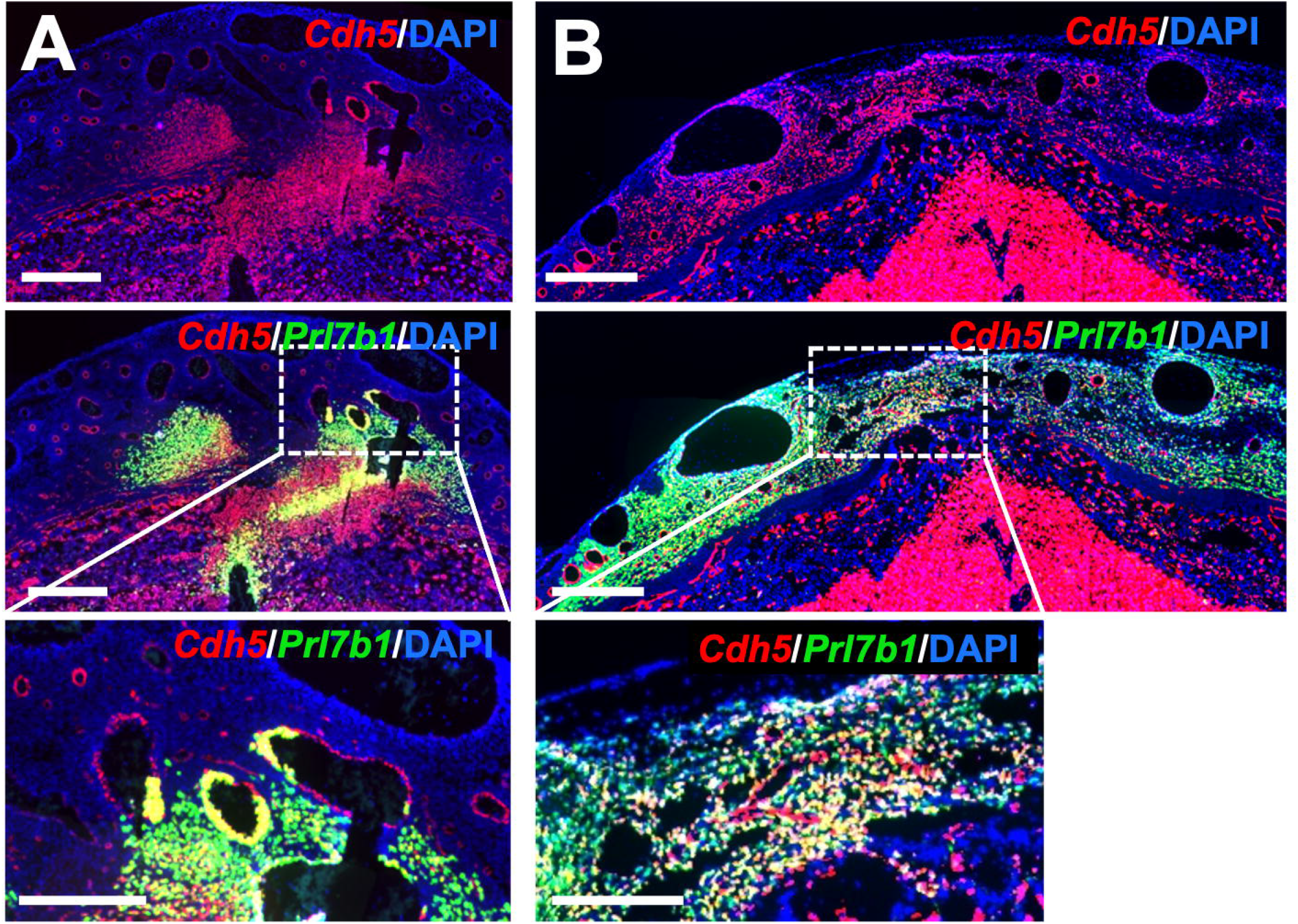
Localization of *Cdh5* in endothelial cells and invasive trophoblast cells. **A**) In situ hybridization localization of transcripts for *Cdh5* (upper) and *Cdh5* and *Prl7b1* (center and lower) within gd 15.5 placentation sites. **B**) In situ hybridization localization of transcripts for *Cdh5* (upper) and *Cdh5* and *Prl7b1* (center and lower) within gd 19.5 placentation sites. The lower panels are high magnification images of regions shown in the central panels. Scale bar=1000 µm.

**Figure S6.**
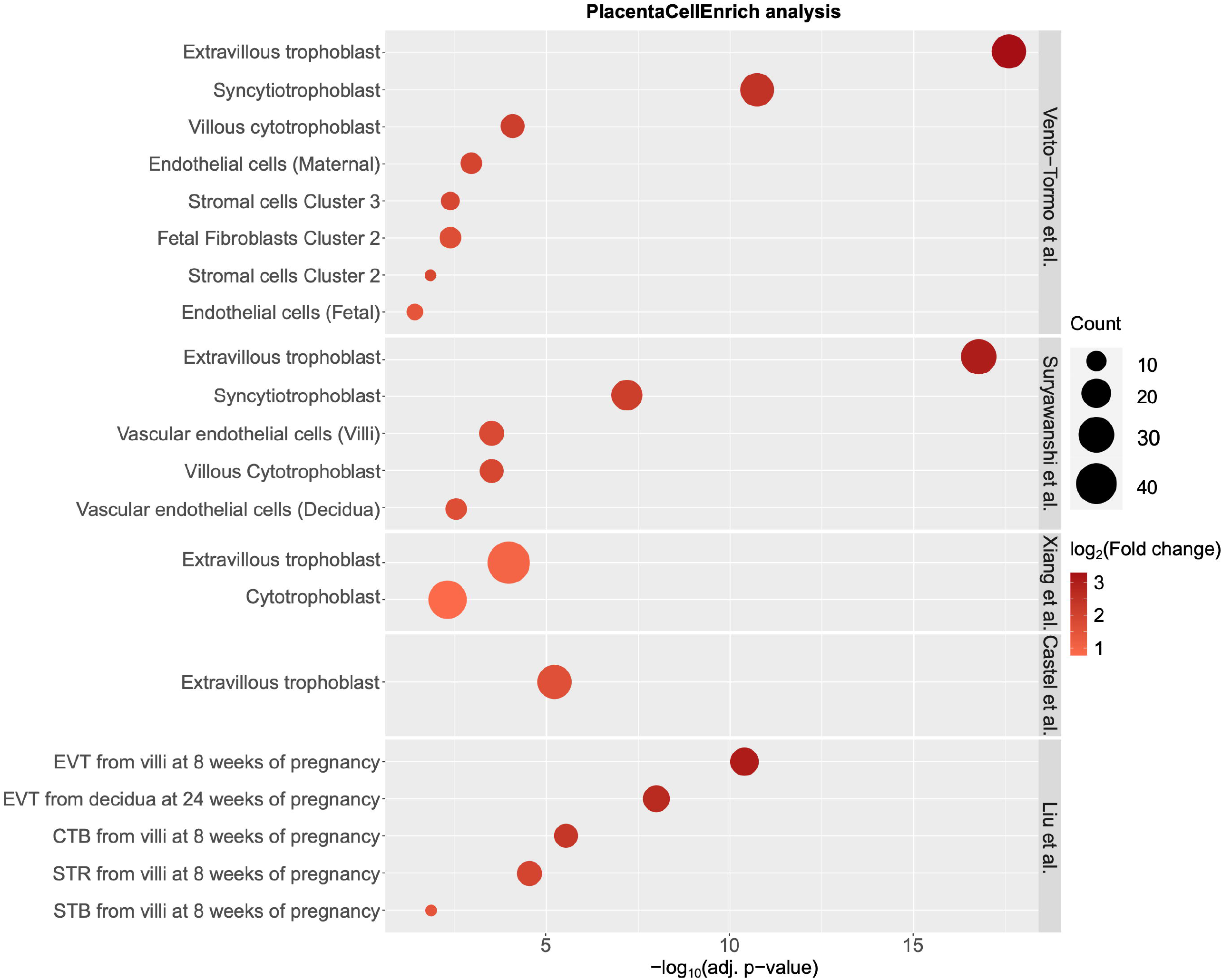
Full PlacentaCellEnrich results. Dot plots showing cell type enrichment when using marker genes from the invasive trophoblast cell gene clusters in rats as input to PlacentaCellEnrich. A significant enrichment has an adjusted p-value ≤ 0.05, fold change ≥ 1.5, and number of observed genes ≥ 5. Dot size represents the number of observed genes specific to a cell type; dot colors correspond to log_2_(Fold change) of the enrichment. Abbreviations: EVT, extravillous trophoblast; SCT, syncytiotrophoblast; CTB, cytotrophoblast; STR, villous stromal cells.

